# ŌvSim: a Simulation of the Population Dynamics of Mammalian Ovarian Follicles

**DOI:** 10.1101/034249

**Authors:** Joshua Johnson, Xin Chen, Xiao Xu, John W. Emerson

## Abstract

No two ovaries are alike, and indeed, the same ovary can change its architecture from day to day. This is because ovarian follicles are present in different numbers, positions, and states of maturation throughout reproductive life. All possible developmental states of follicles can be represented at any time, along with follicles that have committed to death (termed follicle atresia). Static histological and whole-mount imaging approaches allow snapshots of what is occurring within ovaries, but our views of dynamic follicle growth and death have been limited to these tools. We present a simple Markov chain model of the complex mouse ovary, called “ŌvSim”. In the model, follicles can exist in one of three Markov states with stationary probabilities, Hold (growth arrest), Grow, and Die. The probability that individual primordial follicles can growth activate daily, the fraction of granulosa cells that survive as follicles grow, and the probability that individual follicles can commit to atresia daily are user definable parameters. When the probability of daily growth activation is stationary at 0.005, the probability of atresia for all follicles is near 0.1, and the probability of granulosa cell survival is modeled around 0.88, ŌvSim simulates the growth and fate of each of the approximately 3000 postpubertal mouse ovarian follicles in a fashion that approximates actual biological measurements (e.g., follicle counts). ŌvSim thus offers a starting platform to simulate mammalian ovaries and to explore factors that might impact follicle development and global organ function.

**Author Summary:** ŌvSim is a computer simulation of the dynamic growth of mouse ovarian follicles. The program is offered as the beginning of a research and teaching platform to model asynchronous follicle growth and survival or death.

## Introduction

A central goal in reproductive biology and medicine is determining mechanisms that control the fates of mammalian ovarian follicles. This is because follicle growth and survival control the availability of the mature eggs used for conception. Follicles also produce endocrine hormones that are key not only for reproduction, but that support health and quality of life. An ovarian follicle consists of a single oocyte and associated somatic cells. After a period of growth arrest in a ‘primordial’ follicle state, growth activation can occur *via* upregulation of mTOR/Akt signaling (1; 2; 3; 4; 5). Somatic granulosa cells begin to proliferate around the oocyte, which itself grows in size and later resumes and completes meiosis (6; 7). Few follicles survive to the final ovulatory stage where they can release a mature egg; the majority of follicles die within the ovary in a process called atresia (8; 9; 10). Because follicles are present in the thousands in reproductive-age mice and humans, and their growth, development and death occur in an asynchronous, stochastic fashion, it can be difficult to conceptualize the ovary’s function(s) as an endocrine organ and how it achieves its consistent production of mature eggs.

The most common approaches used to account for the developmental states and survival (or death) of mammalian follicles over time is the preparation of static histological sections of ovaries. These are referred to as histomorphometric approaches (8; 11; 12). Histological sections allow a very detailed micron-scale appreciation for all of the cell types and structures in and around follicles. More recently, whole-mount fluorescence analysis has been used to great effect, providing a finely-grained accounting of the numbers and sizes of follicle-enclosed oocytes in the mouse ovary (13; 14). Future modifications of this latter approach may eventually allow for computer-assisted analysis of the disposition of the somatic cells of follicles as well. The primary drawback of static histomorphometric approaches is the need to prepare specimens from many replicate animals at different time points if differences in follicle composition over time are to be appreciated. Experience with this highly laborious process led us to question whether an *in silico* approach of simulation and analysis of follicle numbers over time was possible.

Computer simulations of cells, tissues, and organs are becoming more commonplace. With regards to the ovary, Skodras and Marcelli (15) have produced an interesting graphical and numerical simulation of the size distribution of ovarian follicles in newborn mouse ovaries. Beyond striking graphical images, their study allows the comparison of follicle number in actual (biological) newborn ovaries with realistic simulated counterpart ovaries. As those authors say, such simulations can also support the ”…[analysis of the ovary and other] organs made up of large numbers of individual functional units.” However, tools for the simulation and visualization of dynamic follicle development within the mammalian ovary over the entire reproductive lifespan have not been available. We hypothesized that establishing a simple set of rules for i) follicle growth activation, ii) granulosa cell proliferation, iii) granulosa cell death, and iv) individual follicle survival could provide the necessary starting points for a rudimentary simulation of stochastic follicle behavior over time. Consideration of these rules led us to a Markov chain approach

We reasoned that follicles can exist as growth arrested primordial follicles (a “Hold” state), growing follicles (a “Grow” state), and follicles that have committed to die *via* atresia (a “Die” state). Markov state transition models have been applied as powerful tools in the health and medical literature (e.g., in disease models) (16; 17; 18; 19; 20), and initial modeling of follicles in this way proved fruitful.

We have now produced a function in the R language (21), ŌvSim, to model follicle Markov state transitions across a discrete time series. ŌvSim simulates follicle development and population dynamics according to user-specified starting population of follicles and transition probabilities. To our surprise, the simple probability model, with informally selected and reasonable parameter values, can produce remarkably accurate representations of follicle population dynamics, closely matching the biologically observed number of surviving follicles (and thus an estimate of ovulated eggs) over time. Although this does not prove that the apparently complex process of follicle population dynamics is simple, the results show that a relatively simple probability-dependent process is consistent with and could help us better understand the process of follicle development in nature.

## Materials and Methods

### Ethics statement

“Wet lab” histomorphometric quantification of primordial follicles was performed according to the approved Yale IACUC Protocol #2013-11569.

### Markov chain modeling

The term Markov chain, named after Russian mathematician Andrey Markov (1856-1922), refers to a method for representing stochastic processes by dividing them into unique “states” in a chain. To be modeled by a Markov chain, the states must be considered to be behave independently of any past behavior–a characteristic called “memorylessness.” The probability of moving on to any subsequent state thus only depends on the present state. To model ovarian follicle development using a Markov approach, we establish three states of follicle development (growth arrest, growth, and death; Figure 1) as meeting this criterion. Individual follicles begin in the growth arrest state, and the state can either change or stay the same according to random transition probabilities at each step moving forward in time (in the simulation, days).

**Figure 1.**
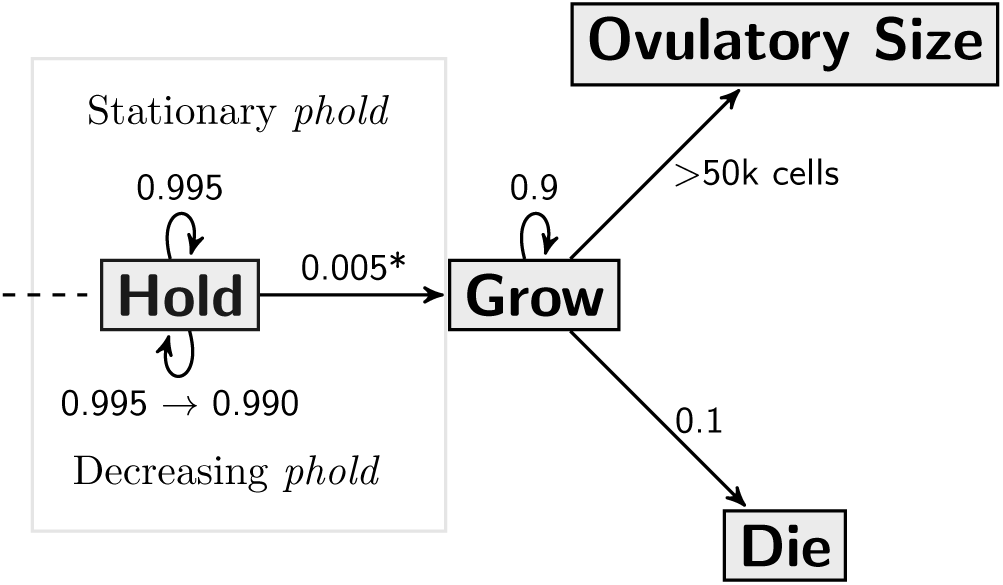
Markov state transition diagram of mouse ovarian follicle development. This flow chart shows the simplified logic of follicle development. From left to right, the first decision for an individual primordial follicle (numerical matrix entry of 1, 2, or 3 granulosa cells) is whether to remain arrested (“Hold”) or to growth activate (“Grow”). This can be simulated as stationary probability *phold* for the duration of the simulation (above dashed line, 0.995), or, as a non-stationary probabillity *phold.new* where the likelihood of remaining growth arrested gradually decreases from 0.995 to 0.990 as the number of growth-arrested follicles reaches a threshold. Growth activation introduces a daily doubling of granulosa cell number. A follicle may then either grow or die daily, with a correction factor of cell death applied to granulosa cell number. If a follicle reaches a threshold number of granulosa cells, it is categorized as an ovulatory follicle.

### Model structure

A simplified example of the Markov matrix operations that we use to simulate follicle growth is seen as follows in (1), where three growth-arrested follicles, *A*, *B*, and *C* are represented by vertical matrix entries populated by one, three, and one “granulosa cell(s).”

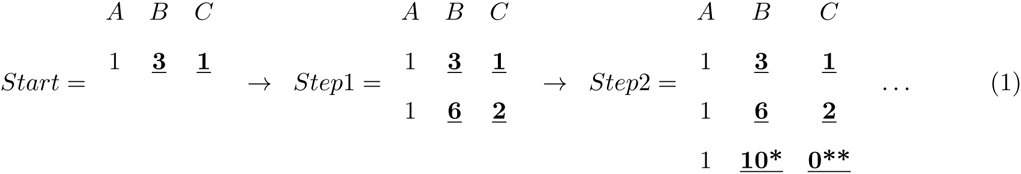

When the simulation begins, follicle states are calculated at each model step according to transition probabilities. In *Step1*, one example follicle (*A*) remains growth-arrested (e.g., its probability calculation results in the “Hold” state) and it maintains its number of pregranulosa cells. The other two follicles (*B* and *C*, indicated by bold, underlined numbers) growth activate in *Step*1 (their probability calculations result in a state change from “Hold” to “Grow”). Existence within the “Grow” state means that follicles can either continue to grow or transition to the “Die” state. While growing, granulosa cell number approximately doubles each daily step. This (daily) doubling time reflects a granulosa cell mitotic index that is consistent with reported (22) and our own (Conca Dioguardi, Uslu, and Johnson, unpublished) data. In *Step*2, follicle *B* grows but is shown to contain less than double the number of granulosa cells in the previous step due to granulosa cell death (*; modeled as a Bernoulli random variable, details in ŌvSim R code, below; after (22) and Conca Dioguardi, Uslu, and Johnson, unpublished). Follicle *C* grew in *Step*1, but commits to atresia in *Step*2 because its probability calculation resulted in the “Die” state, and its granulosa cell number is set to zero (**, see details in ŌvSim R code, below). These steps are represented in the Markov state transition diagram in Figure 1.

### ŌvSim R code and model parameters

ŌvSim R code and accompanying documentation is available on GitHub (https://github.com/johnsonlab/OvSim) and has been released using the MIT License (http://opensource.org/licenses/MIT; see Supporting Information). ŌvSim was designed using known biological parameters of ovarian follicles while allowing users to modify some of these parameters (Table 1). Once the script is activated, a numerical matrix is populated with randomly-generated values corresponding to the starting number of granulosa cells in individual simulated primordial follicles (e.g., one, two, or three pregranulosa cells per (23); see also “puberty” option below).

In ŌvSim, the starting <u>n</u>umber of follicles in the ovary (*NF*), the <u>n</u>umber of <u>d</u>ays of time (*ND*) to run the simulation, and the length of the ovulatory cycle (*cyclength*) can all be specified. We set the number of mouse ovarian follicles to 3000, including 2250 primordial follicles (after (11) and (24)) for most of our studies. Ovulatory cycle length for mice was set at 4, 4.5, or 5 days. As mentioned, we use a daily (e.g., 24 hour) doubling time for granulosa cells and allow users to set the granulosa cell death rate fraction of The script then continues to loop with “daily” probability calculations and operations upon each follicle entry in the matrix. Simulations run for 420 days by default (14 months), corresponding approximately the fertile lifespan of C57Bl/6 mice fed *ad libitum* (25).

Parameters related to follicle growth can be specified as follows. If used, the default *phold* variable is the stationary probability that a primordial follicle stays growth arrested each day. Individual primordial follicles either stay arrested and therefore maintain their cell number of 1, 2, or 3, or, growth activate. Optionally, users can choose to simulate the action of the paracrine factor Anti-Müllerian Hormone (AMH) upon follicle growth activation. AMH produced by growing follicles has been shown to inhibit the growth-activation of primordial follicles (26; 27; 28). When the variable *phold* is set equal to the string *“custom1”*, a non-stationary probability *phold.new* is used in place of *phold*. As the simulation runs, *phold.new* is held at a user-specified value (in our example, 0.995) as follicle numbers decline. When the number of immature follicles declines and reaches the threshold number entered into the variable *threshold*, *phold.new* begins to decline at a user-specified rate per day. In either case, overcoming growth arrest results in growth activation where an individual follicle represented in the matrix is released to a state of exponential granulosa cell growth with a daily doubling time.

Granulosa cell number in growing follicles is controlled by the probability that individual cells within a growing follicle survive (*pcelllive*). Our estimates using histological sections detect a background of pyknotic granulosa cells between 15 and 20% within follicles thought to be intact. Thus (*pcelllive*) is modeled as independent Bernoulli random variable within that range with 0.8 as the default value.

To control the fraction of follicles that commit to atresia, a conditional stationary probability, *cond.pdub* is executed upon each matrix entry each day. As mentioned, the follicle’s matrix entry can double (minus the cell death induced by *pcelllive*, above) with probability *cond.pdub*, Alternatively, the follicle can “die” via atresia with the probability 1 - *cond.pdub*. A follicle’s death is simulated by its matrix entry set to zero.

The parameter *ejectnum* (50,000 by default) reflects the number of granulosa cells required for a follicle to be categorized as a fully mature preovulatory follicle. Critically, the simulation as designed here does not control the final stage(s) of follicle development that ensure that ovulation occurs on only one day per cycle. For now, we are modeling growth patterns that can give rise to approximately ovulatory sized follicles within an entire single ovulatory cycle (4 - 5 days in the mouse). Using (6) as a guide, we set the threshold for survival to ovulation to 50,000 but experimented with thresholds as large as 500,000 granulosa cells.

We also added the ability to optionally begin the simulation placing the ovary in a peripubertal state, where several hundred follicles have already reached the preantral stage of growth awaiting puberty. The option “puberty,” when set to TRUE, populates a user-specified number of matrix entries (variable IGP for <u>i</u>nitial <u>g</u>rowing <u>p</u>ool) with granulosa cell numbers that range from newly growth-activated to the estimated number of granulosa cells in peripubertal preantral follicles. The number of growing follicles and the range of granulosa cells in this prepubertal growing pool can also be user defined.

Overall, it can be said that the model parameters were not formally estimated, but were instead selected based on our domain expertise. The question was whether a simple model of follicle population dynamics might recapitulate apparently complex patterns of follicle growth and survival seen *in vivo*.

### Mice and tissue collection

C57BL/6 mice were handled and tissues were collected in accordance with an active protocol under the auspices of the Yale IACUC. Fresh ovaries were removed and cleaned from the fat, rinsed in PBS and fixed in Dietrich’s fixative (30% Ethanol (EtOH; v/v), 10% Formalin (v/v - using aqueous 37% Formaldehyde solution), 2% Glacial Acetic Acid (v/v); filter prior to use) overnight. Ovaries were then transferred into 70% EtOH for storage at 4°C. Specimens were batched and embedded in paraffin. 5 *µ*m serial sections were cut and placed onto glass slides (Fisher Superfrost/Plus Microscope slides-Precleaned (#12-550-15). Slides were warmed, dewaxed with Xylenes (3 times × 5 min.) and rehydrated through an increasing alcohol series up to distilled water and then PBS. Slides were then stained in Weigert’s Iron Hematoxylin for 10 min. followed by counterstaining in Methyl Blue (0.4 mg/ml in saturated aqueous Picric Acid) for 6 min. Finally, specimens were dehydrated and coverslipped in mounting media (Richard-Allan Scientific Cytoseal-60 Low Viscosity (# 8310-16).

### Histomorphometric follicle counting of primordial follicles

Primordial follicles were counted in every fifth serial section, with raw numbers multiplied by 5 as previously described (29; 30). A follicle was considered primordial if a single layer of flattened pre-granulosa cells surrounded the oocyte.

## Results

### Using ŌvSim to Simulate the Mouse Ovary

To model the development of mouse ovarian follicles over a normal reproductive lifespan, the parameters in the *follicle* function are initialized with “default” values shown in the following function declaration:

~~~
ovsim <- function(NF = 3000,
                ND = 420,
                IGP = 300,
                phold = 0.995,
                cond.pdub = 0.9,
                pcelllive = 0.8,
                cyclength = 4,
                ejectnum = 50000,
                puberty = TRUE,
~~~

Here, 3000 total follicles are present at the start, 2700 of which are primordial (1-3 granulosa cells), and 300 are small growing follicles randomly modeled to have initiated growth in a prepubertal cohort. The estrus cycle length is 4 days, and follicle survival to ovulatory size is “called” if granulosa cell number reaches 50,000. A Markov chain state transition matrix for the stationary probabilities of our three states (Hold, Grow, and Die) according to these settings is shown as follows in (2):

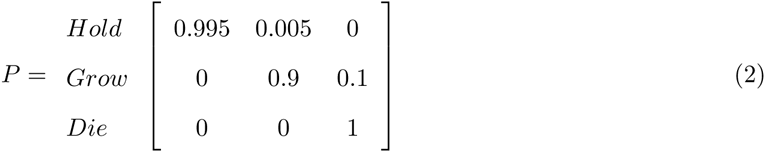

Note that matrix entries that exceed 50,000 simulated granulosa cells are categorized as having survived to ovulatory size, and that follicles that have committed to atresia must stay dead, and therefore their probability of remaining in that state is 1. Figure 1 is a Markov state diagram that includes our default user settings, including the optional setting where the action of AMH upon the probability of primordial follicle growth activation is modeled.

Representative plots of ŌvSim output when an approximately 6-week-old mouse ovary is simulated using default settings are shown in Figure 2. We have expressed model output to highlight how closely ŌvSim resembles key biological ovarian outcomes when the mentioned settings were used. Users can alter model settings or even the code itself in order to test hypotheses about follicle growth and survival.

**Figure 2.**
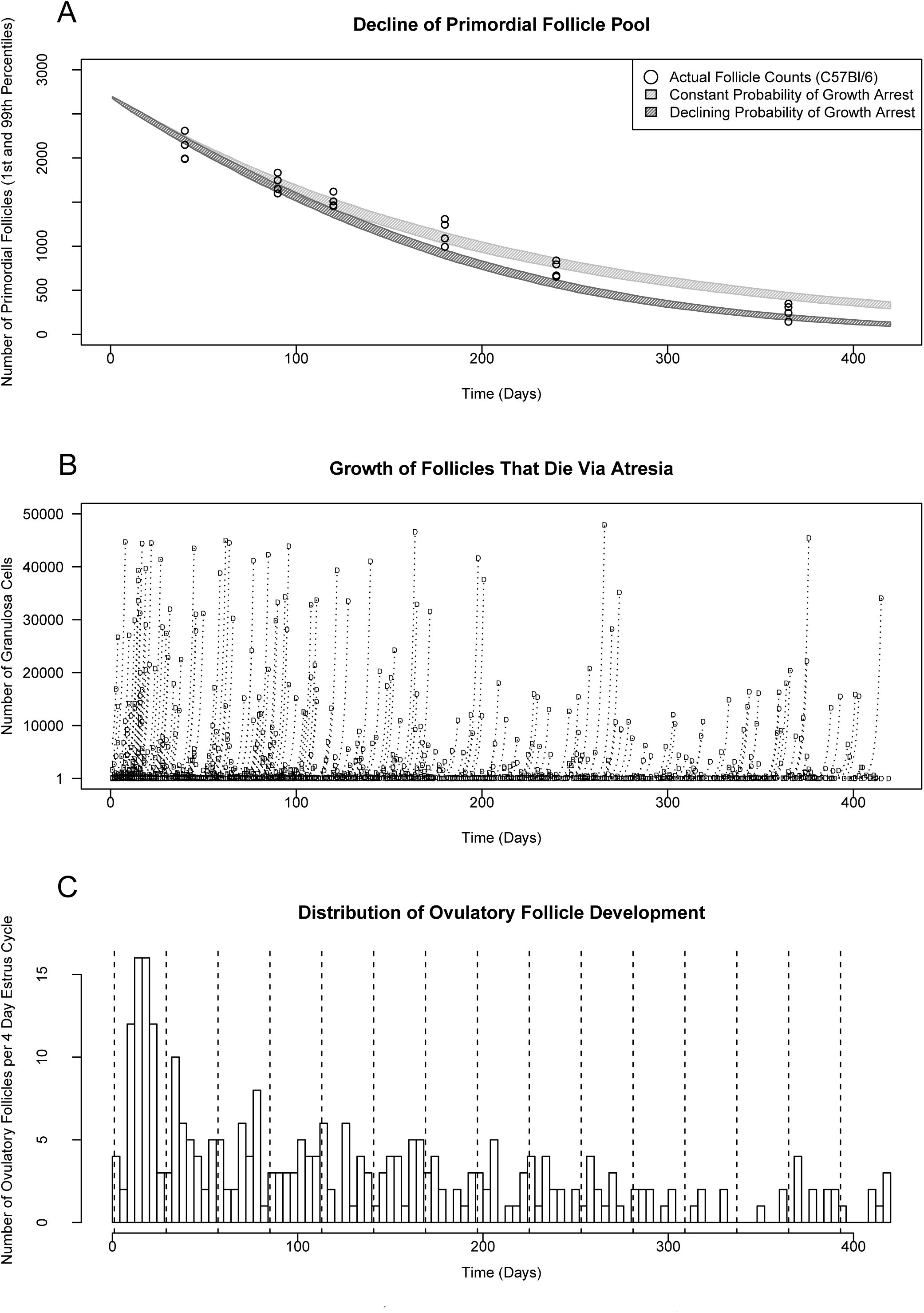
Example ŌvSim mouse ovary output. Default plots produced after an ŌvSim run with “puberty” option set to TRUE, and comparing output for stationary probability of follicle growth activation versus when AMH action is (optionally) simulated. X-axes represent total simulated time in days; pubertal animals would be approximately 50 days old at the start of the simulation. The trajectory of decline of the primordial follicle pool for 1000 ŌvSim runs is plotted as shown in panel **A** where the shaded areas span the first and 99th percentiles of run output when AMH is not simulated (stationary *phold* probability, light gray area) and when AMH is simulated (non-stationary *phold.new*, dark grey area). Circles are actual data from follicle counts of C57Bl/6 mice at 40 days, 3 months, 4 months, 6 months, 8 months, and one year of age (n = 4 ovaries, each from a different animal). In **B**, the growth history of individual follicles that die *via* atresia in a single run (matching run in A when AMH is simulated) is shown by plotting the number of granulosa cells (dashed line) over time, ending with follicle death denoted by the letter ”D.” **C** shows the number of follicles that survive to ovulatory size cutoff (50,000 granulosa cells) each ovulatory cycle (here, 4 days), and dashed vertical lines mark months of time within a single simulation run (*phold.new*, AMH action is simulated).

Panel 2A shows the trend of decline of the primordial pool over time (range between 1st and 99th percentiles) after execution of the simulation 1000 times, comparing the outcome when AMH action is not simulated (gray hatched area, stationary *phold*) versus when AMH action is simulated using a threshold of 100 growing follicles as the trigger for declining probability of growth arrest (black area, non-stationary *phold.new*). Individual data points for actual counts of C57Bl/6 mouse follicles in histological sections at 40 days, 3 months, 4 months, 6 months, 8 months, and one year (circles) are overlaid with simulated data (circles). Panel 2B is a plot of the growth and death of individual follicles that die within the 420 days of simulated time (when AMH action is simulated). Granulosa cell number is represented by the dashed lines, and the time (and follicle “size”) of death is indicated by the letter “D.” Last, Panel 2C is a histogram plot of the distribution of follicles that survive to ovulatory size, grouped in 4 day increments equivalent to the modeled estrus (e.g., ovulatory) cycle length (matches 2B, AMH action is simulated).

The number of eggs available for ovulation each cycle are therefore depicted. ŌvSim also provides CSV-formatted data associated with these plots, useful for finer analyses of follicle size according to granulosa cell number.

### Preliminary application of ŌvSim to human follicle dynamics

Simulation parameters can also be set to conditions mimicking the human ovary, ovulatory cycle length, and approximate reproductive lifespan. For a preliminary human simulation, we set an appropriate number of human primordial follicles at (50,000), the menstrual cycle length to 30 days, and simulated 35 years (unique variable Y, representing the span from approximate ages 15 to 50) of reproductive life. We specified that approximately 1 in 10,000 primordial follicles growth activated per day (phold=0.9999) but kept the follicle survival rate (cond.pdub) and the probability that individual granulosa cells (pcelllive) survive the same as in the mouse simulations (0.88 and 0.75, respectively). For the larger human periovulatory follicle, we set the number of granulosa cells at 500,000. A summary of these parameters as entered follows here.

~~~
human <- function(NF = 50000,
                Y = 35,
                ND = 365*Y,
                IGP = 0,
                phold = 0.9999,
                cond.pdub = 0.88,
                pcelllive = 0.75,
                ejectnum = 500000,
                cyclength = 30,
                puberty = FALSE,
                verbose = TRUE,
                pdfname = NA)
~~~

Representative output from these settings showed follicles that die or survive to ovulatory size in numbers that were reminiscent of human biological outcomes. The total number of follicles that survived to periovulatory size (contain 500,000 granulosa cells) was 605, and the total number of atretic follicles over time was 35403. This meant that 1.4 simulated follicles survived to periovulatory size per month over the length of the simulation. Lacking any additional complexity beyond these parameters, the Markov approach here came very close to the expected “one egg per cycle” output seen in most human natural ovulatory cycles.

## Discussion

The ŌvSim R function simulates ovarian follicle growth using user-definable parameters; when set appropriately, simulations produce results that closely match numbers seen over reproductive life *in vivo*. Follicles that growth activate, die, and reach ovulatory size match the numbers seen *in vivo* over time. Because the R code is freely available for evaluation, use, and alteration, any interested user can contribute to what may eventually become a highly useful simulation of mammalian ovaries. In the meantime, this approach has stimulated interesting discussions about mechanisms that might be at work controlling follicle growth activation, growth, and survival

We emphasize that this is a complete but early-stage simulation using probabilities that account for only a few of the known features of biological follicle development. In the mouse and human ovary, paracrine signaling interactions between follicles impact the rate of follicle growth activation (26) and perhaps follicle survival (31; 32). We and others can work to include finer details of follicle biology in these types of simulations, including the inclusion of additional paracrine and endocrine signaling effects known to affect follicle growth and survival. What is clear here, however, is that stationary probabilities for growth activation and death in our simple model can result in biologically-relevant numbers of follicles that survive to the ovulatory stage or die. The initial inclusion of a non-stationary threshold effect of simulated AMH action resulted in output that matched actual mouse follicle numbers during aging even more closely. As seen in actual mouse and human follicle counts, follicle growth activation accelerates as the number of growing follicles is depleted. While follicle loss is of primary interest, it is also important to consider how synchronization occurs *in vivo* such that those follicles that survive to reach ovulatory size ovulate together on a single day.

The problem of synchronous follicle *availability* for ovulation can be solved by simple rules, but not the more precise follicle *synchronization* such that all periovulatory follicles ovulate on a single day. Selection for ovulation is solved by an additional layer of complexity, the hormonal ovulatory cycle. Ovulation and the final stages of meiotic maturation occur after the LH surge on a single day of the ovulatory cycle, favoring the production of mature eggs on the day of ovulation as well. The hormonal ovulatory cycle can be considered as a “binning” or “winnowing” mechanism, acting upon and selecting follicles of appropriate size to ensure that an appropriate number are ovulated only one day per cycle. The word winnowing can imply the removal of undesirable elements, as in the potential removal of poorer quality oocytes, but whether selection for high quality eggs does occur *in vivo* is unclear (33; 34; 35).

Ensuring that the number of eggs ovulated is tightly regulated in female mammals can be a matter of life or death. Ovulating too few eggs could compromise the survival of a species if too few offspring were produced over time. Ovulating too many eggs can also compromise the survival of a species. Multiple gestation in humans is well known to be a significant risk factor for maternal and offspring loss of life (36; 37; 38). Evolving mechanisms to ensure that the correct number of eggs are produced within an organism’s overall reproductive strategy would therefore have been favored. It is striking that simple regulatory mechanisms (e.g., constant growth activation and atresia rates) can solve much of the problem of the periodic production of ‘safe’ numbers of eggs. Adding a level of ovulatory cyclicity to a future version of ŌvSim will allow the control of ovulation timing to be simulated, and may provide clues about the evolution of the ovulatory cycle itself.

ŌvSim trials show that just a few control parameters can give rise to patterns of asynchronous follicle growth that appear complex. In this initial simulation model, the control parameters only include minimal simulation of interaction(s) between follicles (AMH action). Follicle development within mammalian ovaries may thus in some ways fit the criteria for the phenomenon called emergent behavior (39; 40). Emergent behavior or emergent propert(ies) can appear when a number of simple entities (here, follicles) operate in an environment and form more complex behaviors as a collective (the ovary). Another definition of emergent behavior is any behavior of a system that is not a property of any of the components of that system (40). The mouse (and human) ovary can be modeled as a “system of systems” where overall organ behavior can arise from, but is not necessarily a property of, individual follicles.

Knowing that simple rules can control the number of follicles at different stages of development and death in a fashion that mimics ovarian biology leads to hypotheses that can be tested in ‘wet lab’ experiments. We can test whether similar simple rules underlie ovarian function *in vivo*, and if so, what mechanisms enforce those rules. For example, what mechanisms could control a fixed approximate 1% growth activation rate and 10% overall atresia rate? How can seemingly equivalent primordial follicles growth activate at such a constant rate without activating too quickly or slowly, ensuring that the total duration of ovarian function is appropriate for the reproductive strategy of the female? How can the rate of follicle atresia similarly remain so constant? Premature cessation of ovarian function could result if the rate of either growth activation or atresia were increased (see (41) for a review). ŌvSim can also be modified to model questions that are even more theoretical, such as the impact of ovotoxic agents (e.g., chemotherapeutic or radiological intervention(s)) upon the duration of ovarian function, or, the impact of the additional of new follicles postnatally as suggested by studies that support postnatal oogenesis. The example of the optional modeling of an anti-growth activation factor like AMH highlights the customizable nature of ŌvSim and how users could evaluate the effects of any number of known mechanisms upon simulation output.

The current prevailing consensus in the field is that oogenesis and folliculogenesis ceases before or around birth in most mammals. However, since the first paper calling this into question (24), evidence continues to build that postnatal follicle development can occur *via* the action of female germline stem cells (FGSC) (42; 43; 44; 45; 46; 47; 48; 49; 50; 51; 52; 53). FGSC are currently being used as a source of mitochondria (see (53) for a review) for delivery to oocyte cytoplasm in attempts to improve egg quality and pregnancy rates in the clinic (54; 55). It is a relatively trivial matter to modify the ŌvSim *follicles* function so that new follicles are added at a desired rate and the impact upon the trajectory of follicle loss over time can be estimated. We will continue to develop flexible tools like ŌvSim to address these exciting questions, and hope that other groups will modify the package and build on this approach.

## Supporting Information

### ŌvSim Package Installation

All package and supporting files are available on GitHub (https://github.com/johnsonlab/OvSim) and has been released using the MIT License (http://opensource.org/licenses/MIT). ŌvSim can be installed in an R Environment by following the instructions in the file README.md. Alternatively, the single text file ovsim.R can be executed within R after optional alteration of individual parameters.

### ŌvSim Package License

ŌvSim is available under the conditions of The MIT License (MIT)

© 2015 Joshua Johnson and John W. Emerson

Permission is hereby granted, free of charge, to any person obtaining a copy of this software and associated documentation files (the ”Software”), to deal in the Software without restriction, including without limitation the rights to use, copy, modify, merge, publish, distribute, sublicense, and/or sell copies of the Software, and to permit persons to whom the Software is furnished to do so, subject to the following conditions:

The above copyright notice and this permission notice shall be included in all copies or substantial portions of the Software.

THE SOFTWARE IS PROVIDED ”AS IS”, WITHOUT WARRANTY OF ANY KIND, EXPRESS OR IMPLIED, INCLUDING BUT NOT LIMITED TO THE WARRANTIES OF MERCHANTABILITY, FITNESS FOR A PARTICULAR PURPOSE AND NONINFRINGEMENT. IN NO EVENT SHALL THE AUTHORS OR COPYRIGHT HOLDERS BE LIABLE FOR ANY CLAIM, DAMAGES OR OTHER LIABILITY, WHETHER IN AN ACTION OF CONTRACT, TORT OR OTHERWISE, ARISING FROM, OUT OF OR IN CONNECTION WITH THE SOFTWARE OR THE USE OR OTHER DEALINGS IN THE SOFTWARE.

## Acknowledgments

Drs. Giovanni Cottichio and Taiwo Togun are acknowledged for comments upon the manuscript prior to submission.

